# A semi-automated approach to dense segmentation of 3D white matter electron microscopy

**DOI:** 10.1101/2020.03.19.979393

**Authors:** Michiel Kleinnijenhuis, Errin Johnson, Jeroen Mollink, Saad Jbabdi, Karla L. Miller

**Affiliations:** Oxford Centre for Functional MRI of the Brain, University of Oxford, Oxford, United Kingdom; Sir William Dunn School of Pathology, University of Oxford, Oxford, United Kingdom

**Keywords:** serial blockface scanning electron microscopy, white matter segmentation, corpus callosum, g-ratio, axon diameter distribution

## Abstract

**Purpose:** Neuroscience methods working on widely different scales can complement and inform each other. At the macroscopic scale, magnetic resonance imaging methods that estimate microstructural measures have much to gain from ground truth validation and models based on accurate measurement of that microstructure. We present an approach to generate rich and accurate geometric models of white matter microstructure through dense segmentation of 3D electron microscopy (EM).

**Methods:** Volumetric data of the white matter of the genu of the corpus callosum of the adult mouse brain were acquired using serial blockface scanning electron microscopy (SBF-SEM). A segmentation pipeline was developed to separate the 3D EM data into compartments and individual cellular and subcellular constituents, making use of established tools as well as newly developed algorithms to achieve accurate segmentation of various compartments.

**Results:** The volume was segmented into six compartments comprising myelinated axons (axon, myelin sheath, nodes of Ranvier), oligodendrocytes, blood vessels, mitochondria, and unmyelinated axons. The myelinated axons had an average inner diameter of 0.56 μm and an average outer diameter of 0.87 μm. The diameter of unmyelinated axons was 0.43 μm. A mean g-ratio of 0.61 was found for myelinated axons, but the g-ratio was highly variable between as well as within axons.

**Conclusion:** The approach for segmentation of 3D EM data yielded a dense annotation of a range of white matter compartments that can be interrogated for their properties and used for *in silico* experiments of brain structure. We provide the resulting dense annotation as a resource to the neuroscience community.

## Introduction

The white matter (WM) shapes an integral part of brain function. Not only does it determine the precise long-range anatomical connectivity between brain regions, its microstructure also alters the conduction and timing of physiological signals [1]. To understand the functional architecture of the brain it is required that connections are identified at multiple levels of inquiry [2]. A detailed description of the connectivity of the brain is sought through major collaborative undertakings, such as in the Human Connectome Project [3] using magnetic resonance imaging (MRI). On the most detailed scale, 3D electron microscopy (EM) techniques are used to chart individual connections in simpler animals [4]. Similarly, a large body of research is devoted to a multitude of microstructural properties of individual axons (e.g. myelination [5], microtubules [6], mitochondria [7]). At the macroscopic end, a relatively new field is looking into mapping aggregate indices of WM microstructure in the whole human brain with MRI [8,9].

One of the biggest challenges in neuroscience today is elucidating the links between these scales measured with MRI and EM [10–12]. To interpret the aggregate signal from voxels in MRI, computational models that predict properties from the data are essential. Ideally, these computational models should be firmly grounded in observations at the microstructural level where the signal is generated. Some models have derived their properties from 2D histological data [13], but these data lack the complexity of the 3D structure of the tissue found in a typical MRI voxel. Conversely, 3D EM methods can provide data with a detail that is uniquely appropriate to achieve this goal. The development and validation of MR-based microstructure mapping techniques can benefit greatly from accurate 3D models of WM microstructure. Therefore, we have developed a pipeline for the reconstruction of 3D EM data of the WM that deals with the specific demands of this tissue type.

To render these 3D EM datasets suitable for this purpose, the cellular constituents and features need to be extracted and assessed on their relevance to the influence on the MRI signal. The accurate segmentation of the enormous amounts of data from the 3D EM technique is very challenging, as the size of data precludes manual dense (labelling all voxels) reconstruction approaches for anything but very small volumes. Over the past decades some excellent approaches for segmenting grey matter (GM) have been proposed and published in software libraries. These include manual tracing packages (CATMAID [14], TrakEM [15], KNOSSOS [16]) as well as semi-automated segmentation tools (Ilastik [17], SegEM [18], rhoANA [19]). Ultimately, though, dense connectome mapping also requires inclusion of the long-range, white matter connections. The projectome of the larval zebrafish has already been established with the 3D EM technique using manual tracing [20] and there are efforts to achieve axon tracing throughout the mammalian brain in 3D EM datasets [21]. Nonetheless, comprehensive automated tracing of long-range heavily myelinated pathways currently appears prohibitively demanding, with acquisition times of more than a year for volumes equivalent to a cortical column [22].

While EM-based mammalian connectome mapping is yet out of reach, 3D EM can be used to inform and improve the MRI tools that are commonly employed to assess brain connectivity and microstructure. Our specific goal of segmenting 3D EM volumes is grounded in the general goal of building and validating biophysical models for predicting MRI signals. These segmented volumes can be used as the substrate for such models: for example, simulating the movement of water molecules in tissue for calculating diffusion MRI signals [23] or capturing microgeometry for estimation of microscopic field offsets in susceptibility-based MRI [24]. Monte Carlo diffusion simulations, for instance, have the potential to accurately predict the signals generated in an MRI voxel by a particular diffusion MRI sequences. Realistic simulations require mesh models that include all tissue constituents that influence the movement of water molecules underlying the diffusion MRI signal. This movement is restricted or hindered by cell membranes of unmyelinated axons, the myelin sheath and subcellular structures (mitochondria, microtubules, etc). These have to be modelled as walls in the simulation environment. Ideally, individual objects have to be identified and classified to a particular compartment to assign properties such as permeability and drag that may be very different for various compartments. Therefore, to obtain a comprehensive view of all signal components that contribute to the diffusion MRI signal, dense segmentation of the tissue into individual objects is desired.

Alternatively, highly accurate 3D models could be used to validate conditions under which simplifying assumptions can be made in biophysical models: for example, the use of hollow, impermeable cylinders to mimic axons in diffusion MRI. Finally, these models could provide a platform for in silico experimentation, in which changes to different tissue properties could be explicitly manipulated to predict associated signal changes.

The WM has specific properties that present both challenges and opportunities for dense segmentation that warrant different approaches from those used for GM segmentation. Firstly, while the thick membranous myelin wrappings do greatly facilitate the tracing of myelinated axons, it is not without complications. For instance, Nodes of Ranvier present themselves as gaps in the myelin sheath, necessitating specific algorithms to deal with them. Furthermore, myelin sheaths need to be assigned accurately to individual axons in some applications, including use in diffusion MRI simulations where mesh representations of entire compartments are required. Such segmentations are not always trivial because myelin sheaths often abut each other and oligodendrocytes wrap their processes around multiple axons. A second difference between the two tissue types is the more homogeneous organization of the WM as compared to the GM. As WM is mainly composed of myelinated and unmyelinated axons that are organized in bundles, a few cell body types (oligodendrocytes and astrocytes) and blood vessels, the identification of individual components is simplified compared to GM (where, for example, an important major challenge is the identification of synapses). This relatively simple microstructure makes fully automated segmentation of WM into its cellular components more feasible than for GM.

For the WM specifically, a limited number of automated segmentation approaches have been published. AxonSeg is a Matlab-based library for segmentation of myelinated axons in 2D histology slices [25]. It is primarily based on morphological operations requiring a roughly circular cross section, but has been extended to a more flexible deep learning extension [26]. Kreshuk et al., 2015 [27], leverage the widely used Ilastik tool for segmentation of myelinated axons, complemented with an algorithm to detect and close gaps at the nodes of Ranvier. The recently proposed approach ACSON [28] uses bounded volume growing to provide an axon segmentation of WM tissue from 3D EM datasets, but does not assign the myelin compartment to individual axons. A random walker segmentation has been proposed to segment myelinated axons from 3D-EM volumes for use in an MRI model of orientation dispersion [12].

In this work, we present a pipeline for generating segmentations of WM tissue compartments from 3D EM data that specifically aims to be useful for biophysical models of MRI signals. In particular, we have designed this pipeline for the stringent requirements of realistic Monte Carlo simulations of the diffusion MRI signal using mesh representations of a range of tissue compartments. It builds on well-established open-source tools that have proven accuracy for GM segmentation, but focuses on the unique characteristics of WM tissue. The raw datasets that were used to test the pipeline, the final segmentations, as well as the code are made available. The Python code can be downloaded from https://github.com/michielkleinnijenhuis/EM.

## Methods

### Tissue handling

Two animals were used to collect the data presented here. For the first dataset (DS1), a male adult Balb/c mouse was perfused with Ringer’s solution with 20 units/ml of heparin followed by a mixture of 2.5% glutaraldehyde and 2.0% formaldehyde in 0.1M PIPES buffer and post-fixed overnight in the same solution. The brain was removed and placed in buffer for 48 hours and then bisected mid-sagittally and sectioned at 100 μm using a vibratome. The genu of the corpus callosum was cut from the second full section and this sample was prepared for 3D EM according to the protocol described in [29], except that the 50% resin infiltration step was increased to overnight and the samples were given an extra 48 hrs in 100% resin with multiple changes of fresh resin over this time.

The second animal (dataset DS2) was a male adult MyRF transgenic mouse (not activated). The perfusion of the animal was as above, but used 0.1M sodium cacodylate with 4.35% sucrose as buffer. Vibratome sectioning was done at 300 μm. The corpus callosum was cut from a midsagittal section and was then bisected anterior-posteriorly through the midbody. The anterior sample was prepared for 3D EM up to the dehydration stage as for [29], then the dehydration and Durcupan resin infiltration was performed with microwave assistance, using a Leica AMW.

After EM preparation, the two resin-embedded samples were trimmed to ~0.5×0.5 mm blocks containing the genu, mounted on 3View pins using conductive epoxy and baked at 60 °C overnight. The samples were then coated with ~15nm gold using a Quorum 150 RES sputter coater.

### Electron microscopy data acquisition

Our pipeline () was tested on serial blockface scanning electron microscopy (SBF-SEM) datasets acquired from the corpus callosum of the mouse brain. In SBF-SEM, an SEM image of the blockface is acquired using the backscattered electron signal after which a thin section is removed from the top of the blockface using a diamond knife. This process is automated and repeated many times to build up the high resolution 3D volume with minimal deformations from section to section (for a detailed review of volume EM see [30]). The system consisted of a Zeiss Merlin Compact VP Scanning Electron Microscope (Carl Zeiss Ltd., Cambridge, UK) equipped with Gatan 3View 2XP module.

The first dataset (DS1) was collected with an accelerating voltage of 5 kV in variable pressure mode (50 Pa) using a 30 μm aperture. Images were acquired as a 2×2 montage with 10% overlap each with a frame size of 4000×4000 pixels. The in-plane resolution was 7.3×7.3 nm with a pixel dwell time of 3 μs. The number of sections was 460 with a thickness of 50 nm. This yielded a field of view of ~60×60×23 μm after stitching. The second dataset (DS2) was collected with an accelerating voltage of 3 kV in variable pressure mode (35 Pa) using a 30 μm aperture. Images were acquired with a frame size of 8000×8000 pixels, a resolution of 7.0×7.0 nm (pixel dwell time 4 μs). For DS2, 184 sections with a thickness of 100 nm were collected, yielding a field of view of ~56×56×18.4 μm. Both DS1 and DS2 were taken from the central region of the genu of the corpus callosum, imaged along the sagittal plane.

### Registration

Drift during SBEM acquisition means that slices require slight correction for alignment. Slicewise linear registration was performed using the ‘Register Virtual Stack’ [31] plugin in Fiji [32] using the middle section of the stack as the unmoving reference (maxOctavesize=1024, no shrinking constraint, minimal inlier ratio=0.05). For the montage acquisition, stitching of the sections was performed using the ‘Grid Collection’ plugin [33] (regression threshold=0.30, max/avg displacement threshold=2.5, abs displacement threshold=3.5) with linear blending and subpixel accuracy enabled.

### Pixel classification

A classifier was trained for each dataset using the Ilastik [34] (v1.2.2-post1) ‘Pixel classification’ workflow to assign probabilities (Figure 1: panel3) to each pixel to belong to eight classes (Table 1a). Five of these classes represent compartments of the tissue (myelin, myelinated axons, membranes, unmyelinated axons, mitochondria), while three are annotated to detect the boundaries of the myelin sheaths (inner and outer boundary) and mitochondria compartments (outer boundary). The classes were interactively annotated in a block of 500×500×N_z_ in a minimum of 3 sections (Table 1a: fourth column; 25min/section), continuing annotation for the mitochondria in a minimum of 6 additional sections (Table 1a: fifth column; 2min/section); with sections distributed throughout the block. A subset of the available features was selected to reduce computational load (Table 1b) guided by the ‘Suggest Features’ widget in Ilastik. The classifier was then applied to the full 3D volume.

**Figure 1.**
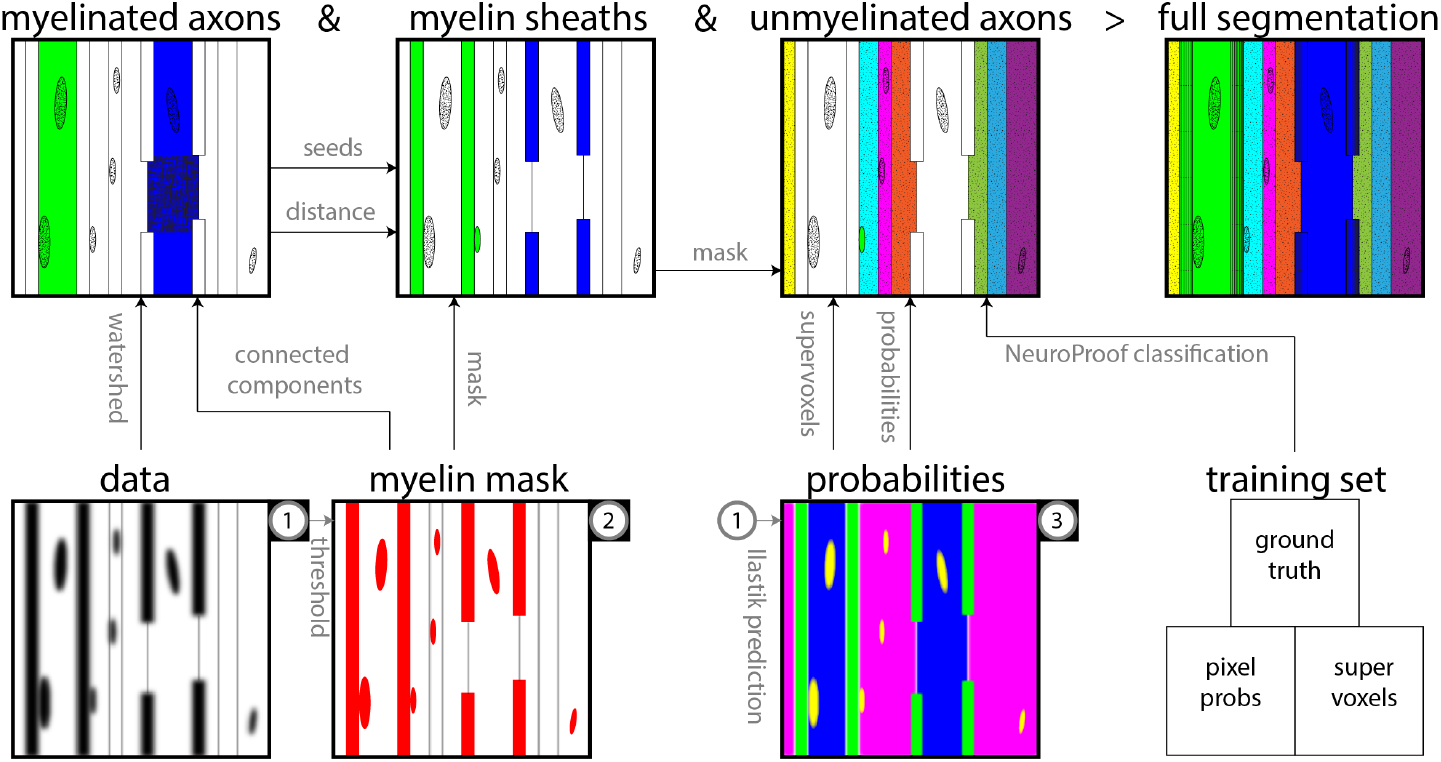
Segmentation pipeline for white matter 3D EM data. General overview of the stages and main relations of the axon segmentation. The full segmentation is formed by consecutively segmenting myelinated axons, myelin and unmyelinated axons (top row). The bottom row lists the minimally preprocessed input volumes for these processing stages (data, myelin mask and probabilities) where arrows indicates which volumes feed into which stages.

**Table 1.**
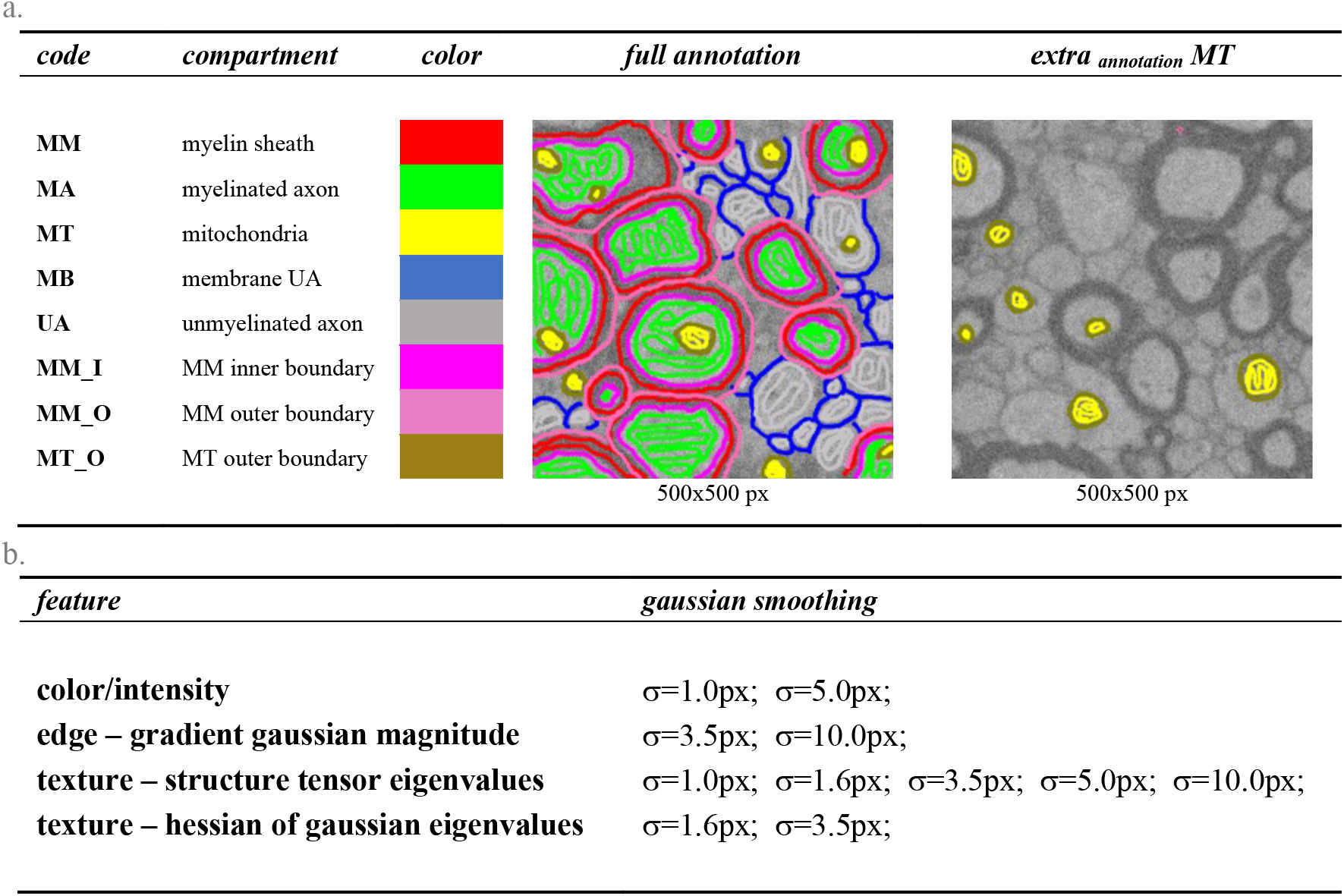
Ilastik pixel classification. a.) Compartments and annotation. b.) Features used in classification.

| *code* | *compartment* | *color* | *full annotation* | *extra annotation MT* |
| --- | --- | --- | --- | --- |
| MM | myelin sheath |  |  |  |
| MA | myelinated axon |  |  |  |
| MT | mitochondria |  |  |  |
| MB | membrane UA |  |  |  |
| UA | unmyelinated axon |  |  |  |
| MM_I | MM inner boundary |  |  |  |
| MM_O | MM outer boundary |  |  |  |
| MT_O | MT outer boundary |  |  |  |
|  |  |  | 500x500 px | 500x500 px |

b.
| *feature* | *gaussian smoothing* |
| --- | --- |
| color/intensity | $\sigma=1.0\text{px}$; $\sigma=5.0\text{px}$; |
| edge – gradient gaussian magnitude | $\sigma=3.5\text{px}$; $\sigma=10.0\text{px}$; |
| texture – structure tensor eigenvalues | $\sigma=1.0\text{px}$; $\sigma=1.6\text{px}$; $\sigma=3.5\text{px}$; $\sigma=5.0\text{px}$; $\sigma=10.0\text{px}$; |
| texture – hessian of gaussian eigenvalues | $\sigma=1.6\text{px}$; $\sigma=3.5\text{px}$; |

### Myelinated axons

Myelinated axons were segmented through a combined 3D/2D connected components procedure using scikit-image [35] (v0.13). First, a myelin mask was created by thresholding the data after smoothing with a 40 nm isotropic gaussian kernel (Figure 1: panel1-2). Small unconnected segments in the otherwise fully connected myelin were removed from the mask by rejecting segments <1.2 μm^3^ (mostly mitochondria). As the in-plane resolution of the sections exceeds the required resolution to detect the myelin sheaths, the myelin mask was downsampled in-plane by a factor of 7 before further processing, taking the 7×7 blockwise maximum for a ~50×50 nm in-plane resolution.

A 3D connected component labelling was performed on the inverse of the 3D myelin mask to segment the non-myelin space (Figure 2: panel4-5). All the connected components of the non-myelin space were labelled in 3D, removing the largest label (representing *un*myelinated axon space) as well as small labels (<0.12 μm^3^; representing small volumes enclosed in between myelinated axons). Erroneous labels were removed in a manual proofreading step using annotation in ITK-SNAP [36].

**Figure 2.**
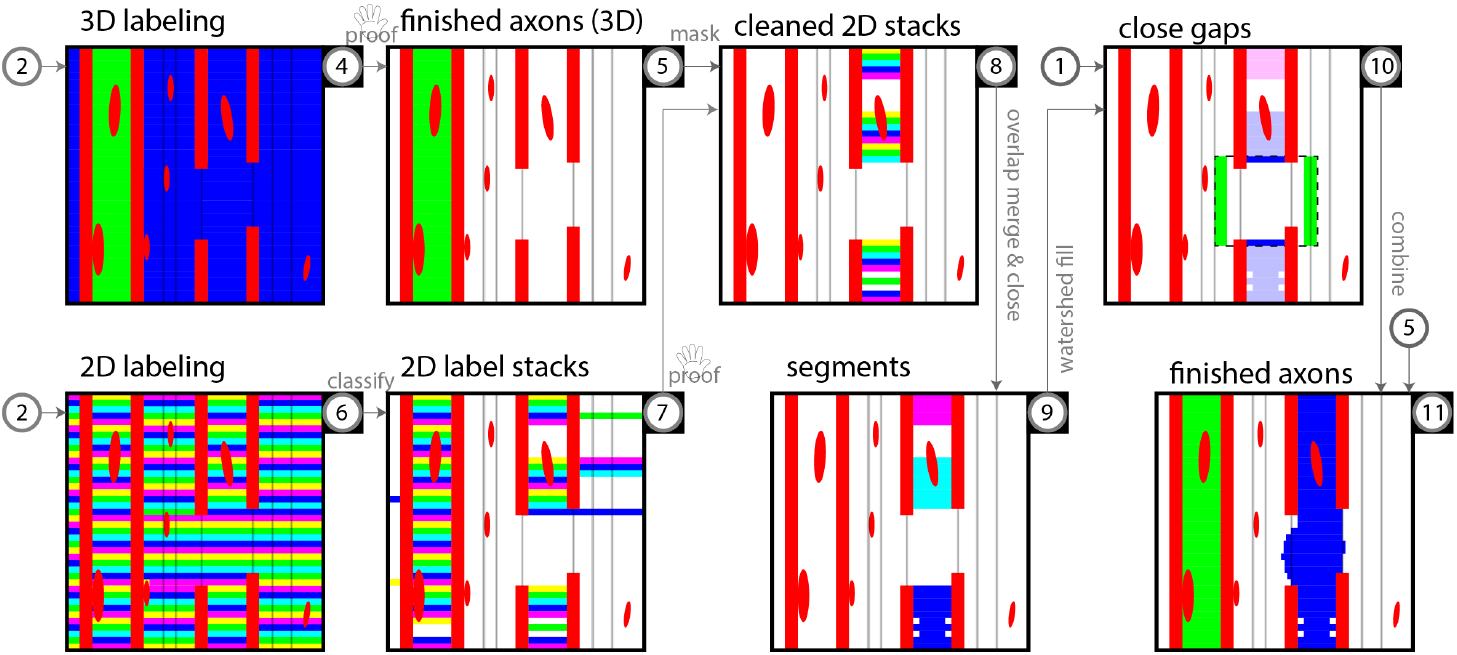
Myelinated axon segmentation. Connected component labeling of the inverse of the myelin mask at low resolution is performed in 3D (panel4-5) and 2D (panel6-11) for non-leaky and leaky axons, respectively. For 3D-labeling, connected components are extracted from the myelin mask (panel4) after which rejection of the largest label and very small labels and manual proofreading yields the non-leaky axons in the volume (panel5). For 2D-labeling, steps can be summarized as 2D connected component labelling (panel6); label rejection by classification (panel7) and proofreading (panel8); merging the label stacks by overlap in neighbouring sections and filling minor gaps (panel9); closing larger gaps by watershed fill (panel10) in a search region (box). The two streams are then combined to arrive at the volume with segmented myelinated axons (panel11).

Because the myelin mask around many myelinated axons does not perfectly enclose the axons, these axons are missed by the 3D labelling. Therefore, a 2D connected component labelling was performed on each z-section (Figure 2: panel6-11) to segment the non-myelin space. Features (Table 2) were computed for each label in order to distinguish labels representing myelinated axons from the remaining space (unmyelinated axons, blood vessels, cell bodies).

**Table 2.** 2D label features.

| *Feature* | *Description* |
| --- | --- |
| **area** | the area of the label |
| **eccentricity** | of ellipse with same second-moments as the label |
| **mean intensity** | of $P_{MA}+P_{UA}$ within the label |
| **solidity** | area / area<sub>convexhull</sub> of the label |
| **extent** | area / area<sub>boundingbox</sub> of the label |
| **euler number** | the euler number of the label |

For the first dataset that was processed (DS1), the selection of myelinated axons was achieved by retaining only labels of which: 1.) the area was between 0.025 and 3.75 μm^2^; 2.) the solidity was > 0.50; and 3.) the extent was > 0.30. Manual proofreading was performed to remove false positive 2D-labels. On DS1 this required extensive manual proofreading. This segmentation served as ground truth to train a support vector classifier in scikit-learn [37] (v0.19.1) based on the features given in Table 2. This more automated segmentation pipeline was then applied to dataset DS2. The classifier was used to predict membership of the myelinated axon compartment for each 2D-label of dataset DS2. As an extra selection step, 2D-labels in which the mean intensity of the myelinated axon probability map from Ilastik classification was larger than 0.8 were all included and 2D-labels with an area larger than 7.5 μm^2^ were all excluded.

After the proofreading step, 2D-labels were aggregated to stacks over the z-direction. Labels that overlap segments identified in the 3D labelling step were first masked from the 2D-labeled volume (Figure 2: panel8). 2D-labels from neighbouring sections were merged according to a criterion of a 50 % overlap. To close minor 1- or 2-section gaps (due to missing labels in the stack), a morphological closing operation is used along the z-direction, after which another aggregation is run, merging newly connected labels using a 20% overlap criterion (Figure 2: panel9).

Any remaining unfinished segments from the 2D-labeled and 3D-labeled volumes (e.g. separated by a node of Ranvier or a series of more than two false negatives in the 2D-labeled volume) were merged and connected through a watershed procedure (Figure 2: panel10). For each segment, merge candidates were sought in a region of 20×20×N_z_ voxels above/below the segment (i.e. positioned above/below the centroid of the 2D-label in the top/bottom section of the segment), where N_z_ was increased in successive iterations N_z_=[10, 40, 80]. In this search region, seeds were placed in the border section: positive seeds consisted of the segment’s 2D-label and the remainder of the voxels in this section were negative seeds, while the myelin space was masked from the watershed operation. Merge candidates were identified by selecting segments that 1) showed an overlap of more than 10 voxels with the positive label after watershed; and 2) did not occupy any of the same sections as the seed segment (i.e. did not backtrack). The segment with the largest overlap was selected for merging with the seed segment.

The gap in between the merged pair was then filled by performing a new watershed using both the 2D-labels in the border sections as seeds. This watershed was constrained within a cylindrical region projected between the label centroids in the border sections of each of the two segments with a radius of double the equivalent radius of the largest seed label. The space outside the cylinder was used as a negative seed, while the myelin space was masked. Finally, segments that did not traverse volume at this stage were mostly disconnected by a node of Ranvier where leaving the volume. The procedure was adapted by using watershed fill to the volume boundary instead of to a connecting segment.

To translate the resulting myelinated axons to the full-resolution volume, an oversegmentation (in which the volume is partitioned into supervoxels: segments consisting of multiple voxels likely to belong to the same structure – larger than a voxel, but usually smaller than the axons themselves) was derived from the smoothed data using a watershed in the space outside the myelin mask. Seeds were defined by thresholding the data and labelling connected components (rejecting components smaller than 0.0024 μm^3^). To avoid any gaps between the upsampled axons and the high-resolution myelin mask, the low-resolution myelinated axon labels were upsampled to the full resolution and dilated to halfway the myelin sheath. The myelinated axons in the full resolution were then obtained by merging any labels in the oversegmentation that overlapped with the upsampled myelinated axons. Any segments of supervoxels extending outside the dilated myelinated axons were removed and additionally, the union of the (non-dilated) myelinated axons and the aggregated supervoxels was taken to ensure continuous axons (i.e. also including the nodes of Ranvier). Thus, the dilated and non-dilated myelinated axon labels were the outer and inner bounds on the myelinated axons at full resolution.

### Mitochondria and nodes of ranvier

Many mitochondria are included in the myelin mask and therefore form holes in the myelinated axons. We want to segment these mitochondria as a separate subcellular compartment, remove them from the myelin mask and include them in the myelinated axon compartment. To label these mitochondria, two iterations of morphological image closing (structure element of *xyz*=[29, 29, 5] voxels) and hole-filling are performed on the mask of the myelinated axons. The morphological closing has the added benefit of smoothing the boundary of the myelinated axons, in particular at the nodes of Ranvier where the boundary was determined by the inner boundary of the upsampled myelinated axons, rather than the aggregated supervoxels. However, because this smoothing operation also adds thin sheets of voxels at the inner myelin boundary—where its surface is concave on the scale of the structure element—that do not represent mitochondria, the final mitochondria segmentation is achieved by morphological opening (structure element of *xyz*=[15, 15, 1] voxels) of the difference between the myelinated axon mask after and before closing, i.e. MA_closed_ - MA.

Another subcellular compartment that is segmented are the nodes of Ranvier. Nodes of Ranvier are characterized by the absence of myelin over a short length of the myelinated axon. The resulting gaps in the reconstructed myelinated axons were bridged by the gap-filling procedure as indicated above. Now, we can easily identify the nodes of Ranvier by evaluating each myelinated axon over 2D sections, marking any 2D labels that are not fully enclosed by the myelin mask. A node of Ranvier was defined as a consecutive sequence of 2D labels in the myelinated axons that are not fully enclosed by the myelin compartment and together span a length of >1 μm.

### Myelin sheaths

The myelin mask represents the totality of all myelin sheaths, many of which are abutting. We aim to represent each myelin sheath as a separate object. To separate the individual sheaths, a watershed procedure is used (Figure 3). In order to generate a seed region for each myelinated axon that closely follows the inner boundary of the myelin sheath, the procedure uses the myelinated axon labels where the mitochondria within the myelinated axons are included, but the nodes of Ranvier are removed (Figure 3: panel12). The seeds are obtained by dilating the myelinated axon labels into the myelin mask. For the watershed’s intensity input / landscape, the Euclidean distance transform is used: for each voxel in the myelin mask, the distance to the nearest voxel in any myelinated axon is calculated. The watershed is constrained to voxels in the myelin mask with a maximal distance of 0.35 μm to any myelinated axon.

**Figure 3.**
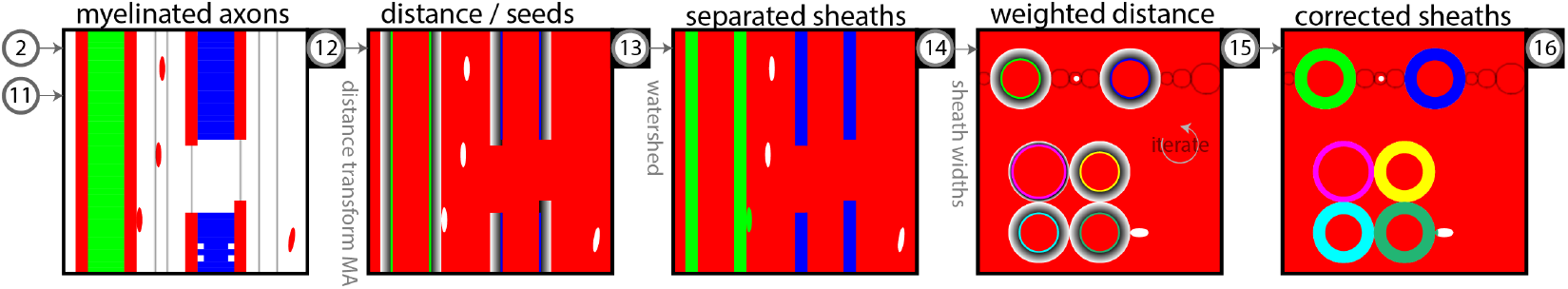
Myelin sheath separation. Subdividing the myelin mask into individual sheaths is based on watershed from the myelinated axon seeds on the distance map of these myelinated axons (panel12-14). Biases in sheath thickness where axons with different thickness touch are corrected through an iterative weighted approach using the axon’s overall sheath thickness in a weighted distance map (panel 15-16). The hand-icon indicates steps where manual proofreading effort is required.

The distance transform is agnostic to the thickness of the myelin sheath of individual myelinated fibres. If two abutting myelinated axons have different sheath thickness, the boundary based on the distance transform watershed will be skewed towards the axon with the thicker sheath. To mitigate this issue, an iterative weighted-watershed is performed, using a modulated distance map. The modulation is derived from the median sheath thickness of the previous pass. The median will be a good approximation of the axon’s thickness under the assumptions that 1) over most of its surface area, the sheath does not touch other sheaths with very different thickness; and 2) the thickness of the sheath is relatively constant over the axon. The weighted distance transform is calculated on a per-axon basis and modulated by a sigmoid function with a width of the median sheath thickness of that axon multiplied by a weighting factor *w* controlling the sensitivity (for this work *w*=10). Per-label weighted distance maps are combined by taken the minimum over all maps. Additionally, in the weighted watershed the mask is constrained to 1.5 times the median width around each myelinated axon (1.2 times for the final iteration).

### Unmyelinated axons, glia & blood vessels

The remaining tissue compartments, mainly unmyelinated axons, are segmented by automated classification using NeuroProof [38], a segmentation method that learns to agglomerate a graph of supervoxels into neurons using features from the provided probability maps (as obtained from the Ilastik pixel classification).

Our supervoxels are generated by watershed of the summed and smoothed (σ=21 nm) probability map for intracellular space (P_ICS_=P_MA_+P_UA_). The seeds are obtained by finding local maxima in the ICS probability map that are >0.1 μm apart and exceed PICS=0.8. We isolate the unmyelinated axon space by masking out the myelinated axons as identified in the previous steps (axons, sheaths, mitochondria and nodes). In addition, we mask out the mitochondria of the unmyelinated axons. These are hypointense in the ICS probability map and we define their mask by thresholding at P_ICS_=0.2.

A ground truth segmentation was generated for a block of 500×500×430 voxels of dataset DS1 by manually proofreading and merging the supervoxels in that block. Next, a random forest classifier is trained on this annotated training dataset with NeuroProof (settings: 5 iterations; strategy type 2; no mitochondria context). Finally, with this classifier, the supervoxels of the full datasets are agglomerated to form the processes of unmyelinated axons and glia, glial bodies and blood vessels with a threshold setting of 0.5. This stage requires extensive proofreading to correct split/merge errors, although we chose not to pursue this here. Conversely, we have improved on the output of the random forest classifier by specifically identifying the large structures in the dataset (the glial bodies and blood vessels; further subdivided into glial bodies, glial processes surrounding bodies, blood vessel lumen, blood vessel walls, pericytes). This was achieved by performing a partial manual annotation of each 10^th^ slice (x-direction) in the low-resolution dataset using ITK-SNAP after which these annotations are upsampled and the supervoxels that overlap with the manual annotations are agglomerated to form these additional compartments.

## Results

Dataset DS2 serves as an example of the detailed workflow and will be used for demonstrating the pipeline’s features and limitations.

### Preprocessing

The 8-class probability map from Ilastik (Figure 4d) suggests that, as expected, the myelin boundaries are well-defined (MM, MM_I, MM_O); the classifier can distinguish between intracellular spaces for myelinated (MA) and unmyelinated axons (UA); the thin membranes of unmyelinated axons (MB) are well-separated from the thick membrane wrappings of the myelin compartment; however, the mitochondria probability maps (MT, MT_O) contain high probability for myelin sheaths as well, indicating the general difficulty of separating the myelin and mitochondria.

**Figure 4.**
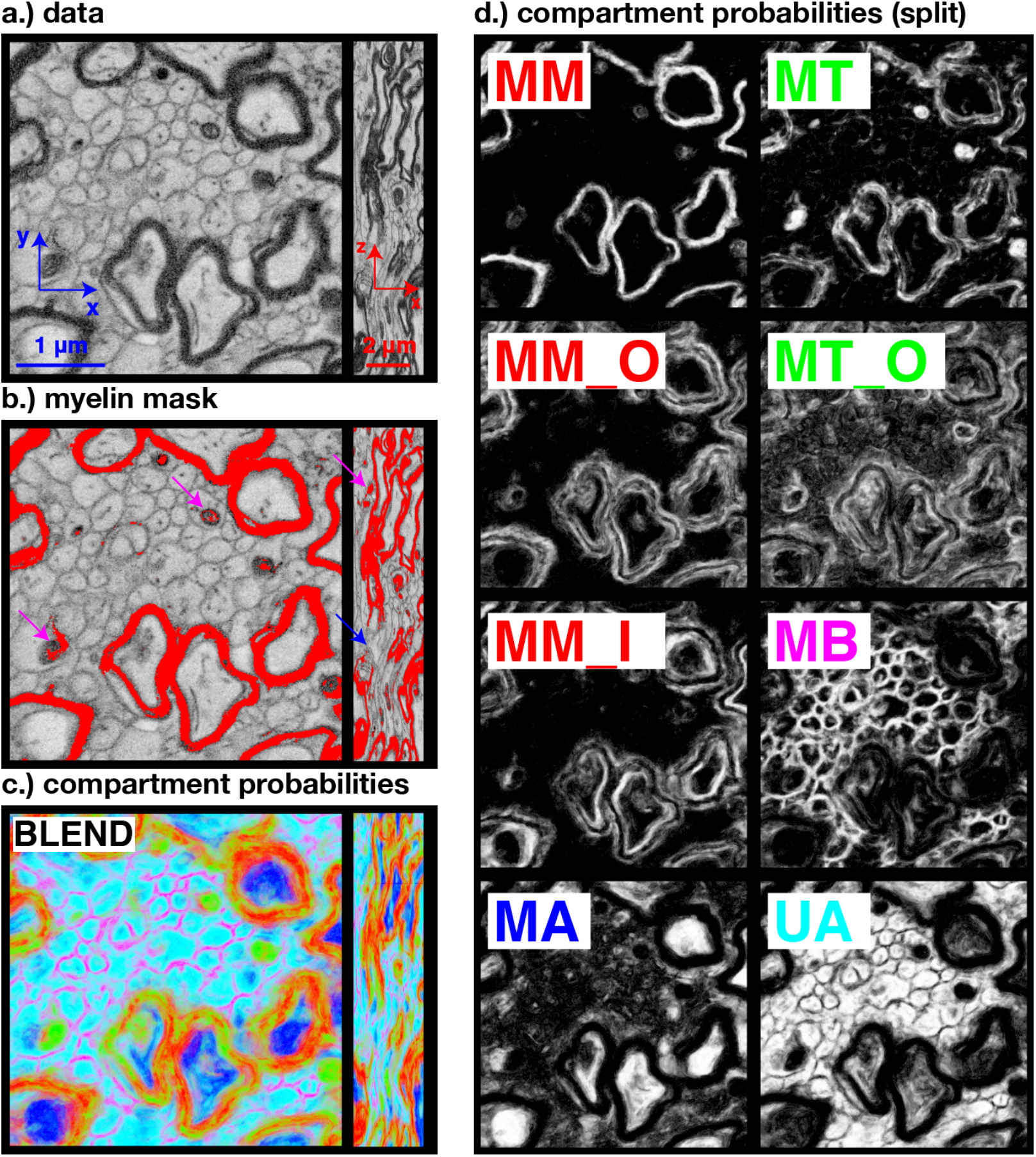
Preprocessing results. a.) Block of 500×500×430 voxels of DS1 after registration. b.) The myelin mask is obtained by thresholding the data after smoothing. The blue array indicates a node of Ranvier; the magenta arrows indicate mitochondria included in the myelin mask. c.) Compartment probabilities from Ilastik pixel classification. For compartments that were split into multiple classes (MM, MT), the colour of the constituent classes are equal. d.) Compartment probabilities split over each class.

### Myelinated axons

The 3D-labelling stage in the segmentation of myelinated axons (Figure 5b) detected 1603 labels, after rejecting the largest label (representing the—almost completely connected—unmyelinated axon space) and labels smaller than 0.12 μm^3^. A further 45 labels were rejected manually (required time: 25 min), because they represented false positives enclosed between clusters of myelinated axons. 1422 labels traversed the volume (Figure 5b: blue labels) and were considered complete myelinated axons. The 136 segments that did not traverse the volume were partial axons, either because the segments were split by mitochondria included in the myelin mask; or because the axon featured a very thin segment disconnecting the segments in the downsampled mask (Figure 5b: green labels; white arrow). These segments are later merged with other segments as part of the 2D-labelling stage.

**Figure 5.**
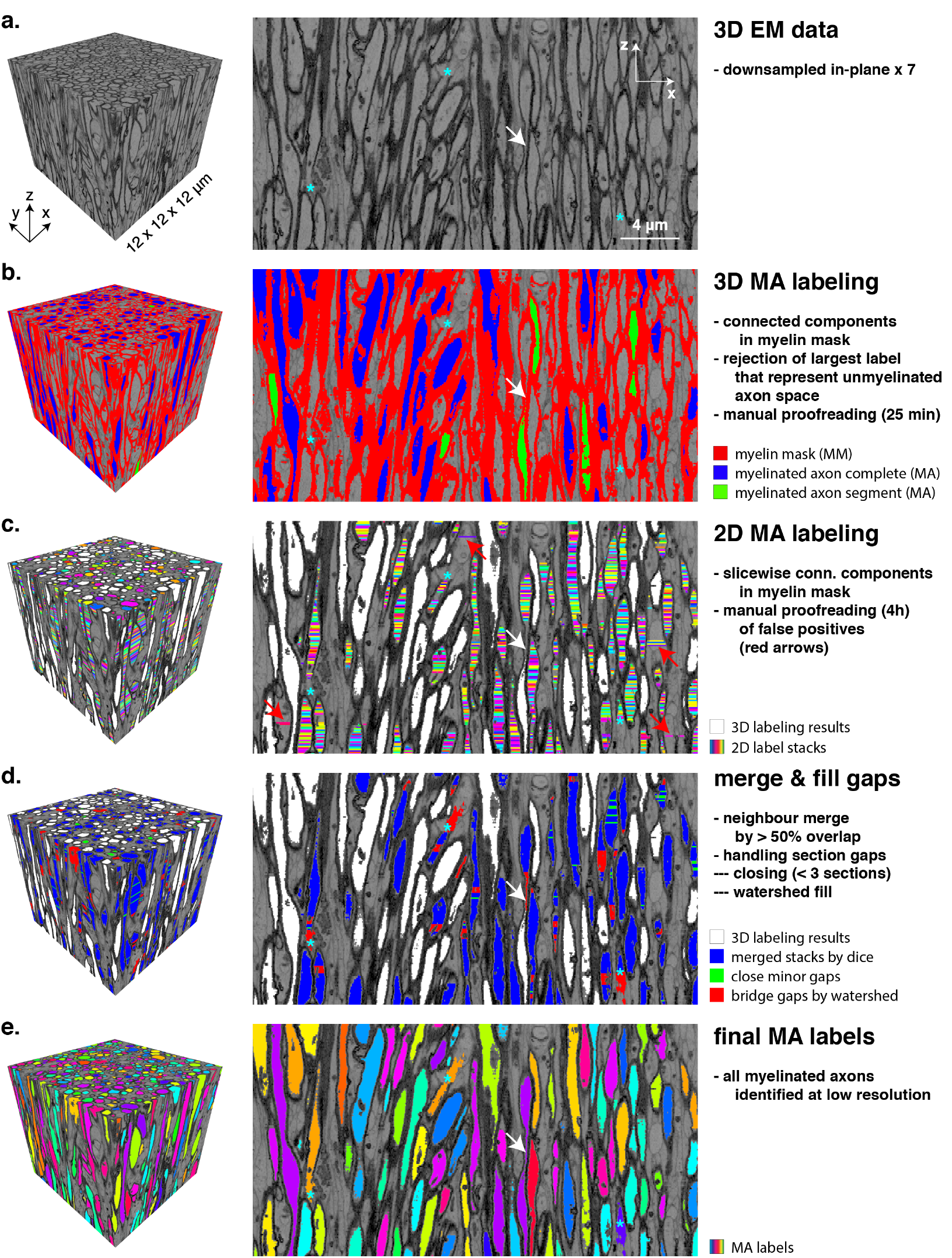
Procedure for identification of myelinated axons (MA). The left and right panels show a small 3D block and a slice view perpendicular to the direction of sectioning, respectively. a.) Data is downsampled in-plane for segmenting the MA compartment. b.) A myelin mask is created by thresholding the data (red). An initial set of myelinated axons is identified by 3D connected component labeling. Labels traversing the volume (blue) were marked as finished, while segments (green) were transferred to the 2D-labeling stage. c.) 2D labels after label classification. Stacks of 2D labels are obtained by slicewise 2D connected-component labeling of the non-myelin space. The space already segmented by the 3D labelling procedure is masked and labels representing myelinated axons are selected by automated classification. Residual false positives (e.g. red arrows) are removed by manual proofreading. d.) Merging neighbouring labels by spatial overlap and closing gaps in between the resulting segments using morphological closing (green; < 2 sections) and watershed fill (red); labels identified by the 3D-labeling stage are masked in white. e.) Final segmentation of myelinated axons at low resolution. Cyan asterisks indicate nodes of Ranvier. The axon indicated by the white arrow (a) exemplifies various steps in the pipeline. It is not fully reconstructed in the 3D-labeling stage (b); it has consecutive false negatives in the 2D-label stacks (c), but it’s segments formed in the 3D- and 2D-labeling stages could be merged by the watershed-fill (d) to a full axon (e).

The presence of nodes of Ranvier and other (unintended) holes in the myelin mask results in leaky myelinated axons, which prompted a 2D-labeling approach for this compartment (Figure 5c). For the first dataset (DS1), 2D-labels belonging to the myelinated axon compartment were filtered according to a set of area and shape criteria, after which a substantial proofreading effort was required to exclude false positive labels outside myelinated axons (required time: ~80 h). The classifier that was trained on the basis of dataset DS1 resulted in a considerably improved initial classification of the 2D-labels of the subsequent dataset DS2. In DS2, both the filtering and classification procedures were applied. The performance of two methods was then evaluated – retrospectively, after proofreading of the myelinated axon compartment to establish the ground truth myelinated axons for DS2. Supplementary Figure 1 compares the the feature-filtering (Fig S1a) and feature-classification (Fig S1b) approaches in terms of label assignment errors. False positives and have significant negative impact on subsequent processing as they create erroneous axons. False negatives result in gaps in the axons and need to be handled by gap-closing and filling procedures. Using the automated classifier rather than filtering, false positives reduced from 138,282 labels to 23,100 labels; and false negatives were increased from 27,902 to 54,363. The percentage of correctly classified labels increased from 81% to 92%. The manual proofreading after classification could now be done in ~4 hours for dataset DS2.

The 2D-labels were further processed by progressively aggregating them into larger segments, until the axon traversed the volume. The first step consists of merging neighbouring labels into stacks (Figure 5d; blue mask) using a criterion of a spatial (Dice) overlap of >50%. Next, gaps of 1 or 2 sections were filled by morphological closing in the z-direction (Figure 5d: green mask). After closing, newly connected label segments were merged through a second run of the overlap merge using a 20% overlap threshold. The result of merging neighbouring 2D-labels and closing small gaps consisted of 179 finished axons, but also of 10,195 label segments that did not traverse the volume.

Thus, the majority of leaky myelinated axons are still fragmented after the first merging and closing attempt. These axons consist of segments with larger gaps, because they have had several consecutive 2D-labels rejected (i.e. false negatives) in the 2D-label classification step. Often, these locations had atypical cross-section (e.g. narrow necks) or represent nodes of Ranvier (indicated by cyan asterisks in Fig. 5) where the 2D-labels flood into the neighbouring unmyelinated axon space.

The watershed procedure to merge and fill these unfinished segments (including those of the 3D labelling stage) was performed iteratively using a progressively larger search region in the z-direction while masking out the finished axons after each iteration. Iterations with closing extents of 10, 40, 80 sections yielded 441, 413 and 51 axons, respectively. The gaps closed by the watershed merge procedure are depicted by the red mask in Figure 5d. Labels smaller than 0.12 μm^3^ voxels were removed at this stage, as they almost always represented residual false positives (missed in the 2D-label proofreading) in the unmyelinated axon space. A final iteration where the myelin mask was not used to constrain the watershed resulted in another 463 axons. This left 871 segments which were not merged by the procedure and were merged by manual intervention (time required: ~16 h). The final myelinated axon segmentation contained 3605 axons (Figure 5e).

Segmentation of the myelinated axons is completed in the full-resolution data. To obtain the myelinated axons at full resolution, an oversegmentation was aggregated by overlap with upsampled myelinated axons (Supplementary Figure 2). The myelinated axon compartment is fine-tuned and subdivided by handling mitochondria and nodes of Ranvier (Figure 6). Mitochondria (Figure 6a) were first included in the myelinated axons by morphological closing of the axons and filling the holes left by the mitochondria. Nodes of Ranvier (Figure 6b) were identified by finding consecutive series of 2D labels (>1 μm along the *z*-direction) that were not enclosed by a myelin sheath. In 3605 myelinated axons, representing a combined length of 27.25 mm, 429 nodes of Ranvier were detected with a median length of 1.9 μm.

**Figure 6.**
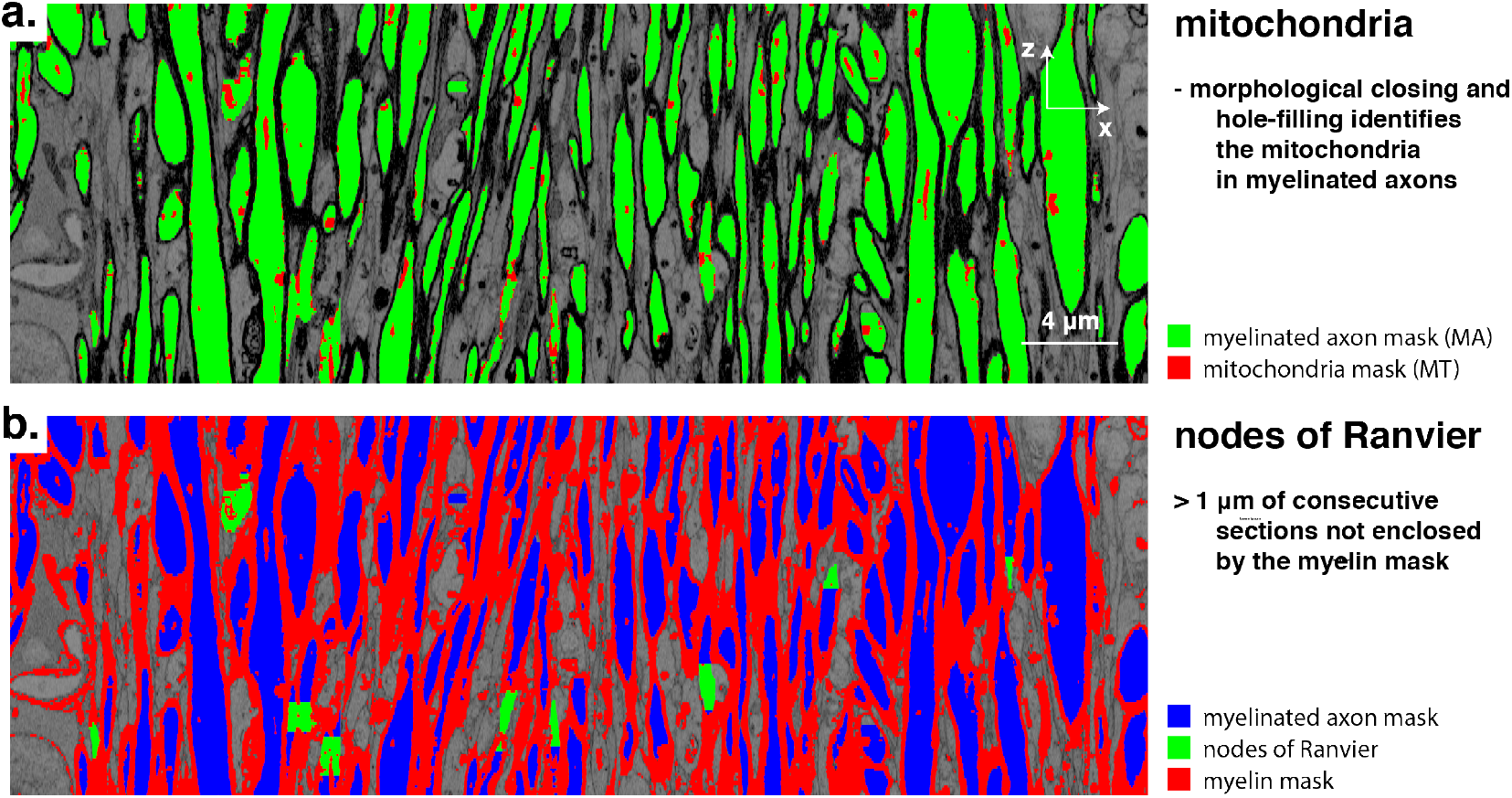
Subdivision of the myelinated axons. a.) mitochondria in myelinated axons (marked in red) are obtained by morphological closing and hole-filling. b.) nodes of Ranvier (green) are interuptions in the myelin (red) around the myelinated axons (blue) and were defined as > 1 μm of consecutive sections not enclosed by the myelin compartment.

### Myelin sheaths

With the myelinated axons and nodes of Ranvier carefully segmented, the identification of myelin sheaths is straightforward. Individual myelin sheaths encapsulating the axons were yielded by subdivision of the myelin mask (Figure 7a; red mask) using a watershed of dilated myelinated axon seeds (Figure 7a; coloured labels) on the distance transform of the myelinated axon compartment (Figure 7b).

**Figure 7.**
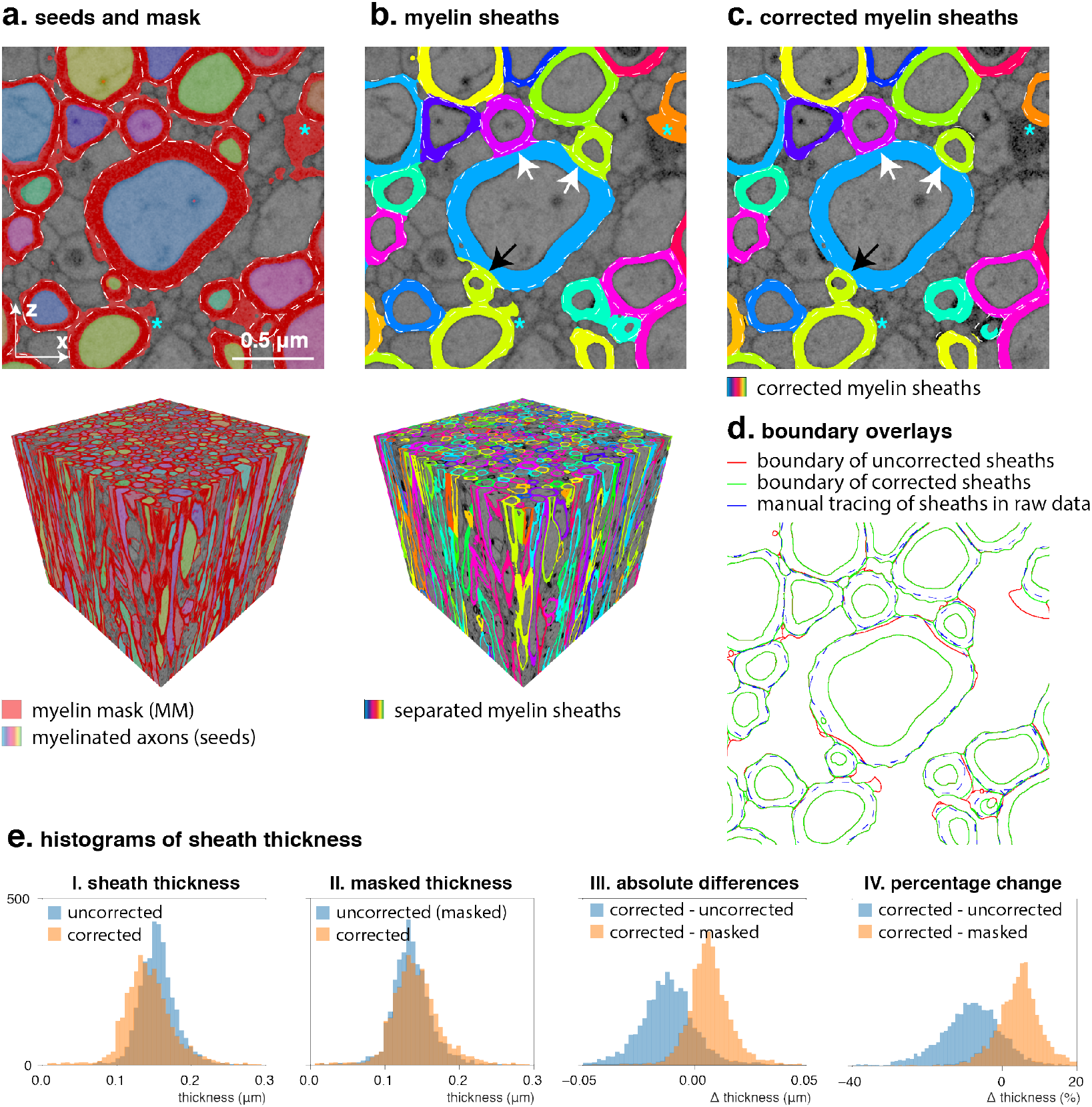
Myelin sheath separation and correction. a.) Myelinated axons are used as seeds to separate the myelin mask (red) into individual sheaths with a watershed on the map of the distance from the myelinated axon compartment mask. Asterisks indicate where mitochondria are included in the myelin mask. A manual tracing of the sheaths is outlined with the white dashed line. b.) Separated sheaths show errors where myelin sheaths of different thickness are touching, leading to overestimation of thin sheaths and underestimation of thick sheaths (arrows). c.) Sheaths after 5 iterations of weighted watershed. d.) The boundary of the corrected sheaths (green) follow the manual tracing (blue) more closely as compared to the boundary of the uncorrected sheaths (red). e.) Corrected sheaths have a lower overall thickness mostly because they are constrained within a mask of 1.2x the sheath thickness which decreases the myelin volume by removing erroneously included (mitochondria) voxels from the sheaths (panel I). The extent of voxel reassignment to different labels can also be evaluated by comparing the thicknesses of corrected vs uncorrected sheaths within the final corrected myelin mask which removes the effect of a difference in total myelin volume (panel II). While the overall thickness decreases, within this mask most sheaths show thickness increases (panel III). The corrections are substantial (panel IV) with a typical 0—30% (mean 9.25%; std 10.9%) decrease in sheath thickness after correction; and thickness changes due to voxel label reassignments typically ranging from −10—20% (mean 4.38%; std 10.0%).

However, the accuracy of this initial segmentation can be poor. One source of error is that, based on this processing pipeline, myelin sheaths include mitochondria of unmyelinated axons (Figure 7a; asterisks). Using an upper limit of 0.35 μm for the myelin sheath thickness partly removes these mitochondria (Figure 7b; asterisks).

A second inaccuracy concerns abutting myelin sheaths with different sheath thickness—a commonly observed configuration. In these locations, the separation of sheaths is skewed towards the thicker sheath (Figure 7b; arrows). In addition to inaccuracies in the estimation of the sheaths’ thickness, the incorrect attribution of voxels of the myelin mask results in distortion in the sheaths’ quasi-cylindrical geometry (e.g. black arrow).

We have attempted to counter these errors through a weighted distance transform to shift the sheaths’ outer boundaries towards their mostly likely true position as derived from the median width over their entire length. Figure 7c shows the individual sheaths after running five iterations of the weighted-distance watershed procedure proposed to mitigate this issue. The restoration of the sheath geometry to circular cross-sections is best appreciated in Figure 7c, while the overlay of boundaries in Figure 7d demonstrates a better overlap between the manually traced boundaries (blue trace) and the corrected sheaths (green trace) as compared to the uncorrected sheaths (red trace).

Beyond the improved separation of abutting sheaths, the individual sheath thickness was used for improvement of the sheaths’ outer perimeter by constraining it within 120% of the median sheath thickness from the myelinated axon. This also improved the exclusion of mitochondria of the unmyelinated axons (Figure 7c; asterisks). In effect, this reduction in the myelin volume by excluding misclassified voxels accounts for most of the difference in sheath thickness distribution between the corrected and uncorrected sheaths (Figure 7e; panel I). Evaluating the sheath thickness distributions within the mask of the corrected sheaths, i.e. removing the effect of changes in total myelin volume and only looking at voxel label reassignments within this mask (Figure 7e; panel II), it is observed that after the correction most sheaths actually increased in thickness. This is obviously at the expense of previously overestimated sheaths that decrease in thickness. Whereas the thickness decreases by an average of 9.3% between the uncorrected and corrected sheaths due to the better-informed distance threshold, within the final mask the median thickness regularization tends to shift the sheaths towards a larger thickness between 0-20% with an average of 4.4% (Figure 7e; panel IV).

### Unmyelinated axons, glia & blood vessels

The space not occupied by myelinated fibres was subdivided into individual unmyelinated axons, blood vessels, cell bodies and cell processes by NeuroProof [38]. The three inputs to train the classifier (probability maps, oversegmentation and ground truth) are shown in the top panel of Figure 8.

**Figure 8.**
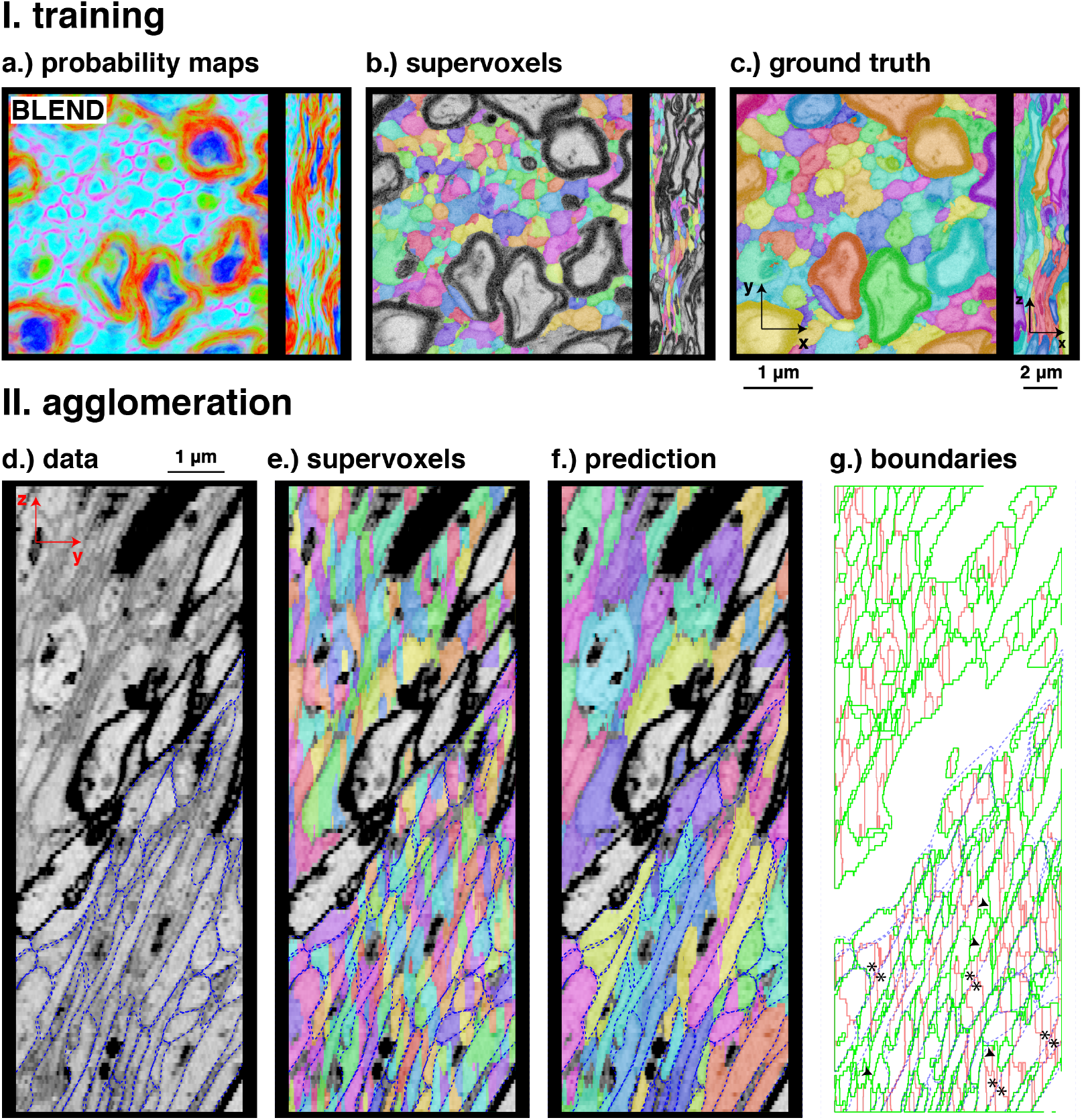
Segmentation of unmyelinated axons with NeuroProof. Panel I: training data. Panel II: example of agglomeration result. a.) A 500×500 section showing probability map outputs of the 8-class Ilastik pixel classifier in a colour blend (red—pooled myelin classes (P_MM_+P_MM_I_+P_MM_O_); green—pooled mitochondria classes (P_MT_+P_MT_O_); magenta—membranes of unmyelinated axons; cyan—unmyelinated axons; blue— myelinated axons). b.) watershed oversegmentation derived from summed intracellular probability maps (P_MA_+P_UA_). c.) ground truth annotation of a block of 500×500×430 voxels. d.) data from a block of dataset DS2, overlaid with manual tracing of unmyelinated axon boundaries (blue dashed lines); e.) watershed oversegmentation; f.) agglomerated axons predicted by NeuroProof; g.) overlay of the manual tracing (blue) and the boundaries of the oversegmentation (red) and agglomeration (green). Although the result in g appears plausible at first glance, a large amount of split (arrowheads) and merge (asterisks) errors remain in the segmentation by fully automated classification without proofreading.

As an indication of the coarseness of the oversegmentation (Figure 8b), the training volume, as masked to exclude the myelinated axons and sheaths, contained 3190 supervoxels, while the unmyelinated axon count in the ground truth was 164 (Figure 8c).

The bottom panel of Figure 8 shows an example of prediction of unmyelinated axons (Figure 8f) and compares it to a manual tracing of the axon boundaries (Figure 8d). The boundary overlay in Figure 8g indicates that although most merges (that have occurred in location where the red boundaries are visible) are correct, many supervoxels remain separated that should be merged (e.g. arrowheads) and some supervoxels have been merged erroneously (asterisks).

### Compartment properties

In this section, we use the final segmentation of the SBF-SEM volume to demonstrate the ability to extract estimates of compartmental properties of biological relevance. Properties of WM tissue that have received considerable attention in the MRI community are the axon diameter, the g-ratio (the ratio of the inner and outer diameter of the myelin sheath) and orientation dispersion. These properties relate to the fundamental function of WM and are of interest in health and disease. One area of active research is to estimate these, and related, microstructural properties using advanced MRI acquisition methods. However, these estimates remain controversial due to the need for strong assumptions in the associated biophysical models and the difficulty in sensitising the signal to these properties with conventional MRI scanners. As comparison between these MRI estimates and EM in the same tissue would be of particular interest, Figure 9 shows examples of some of these properties varying along the length of 200 randomly selected myelinated axons of dataset DS2.

**Figure 9.**
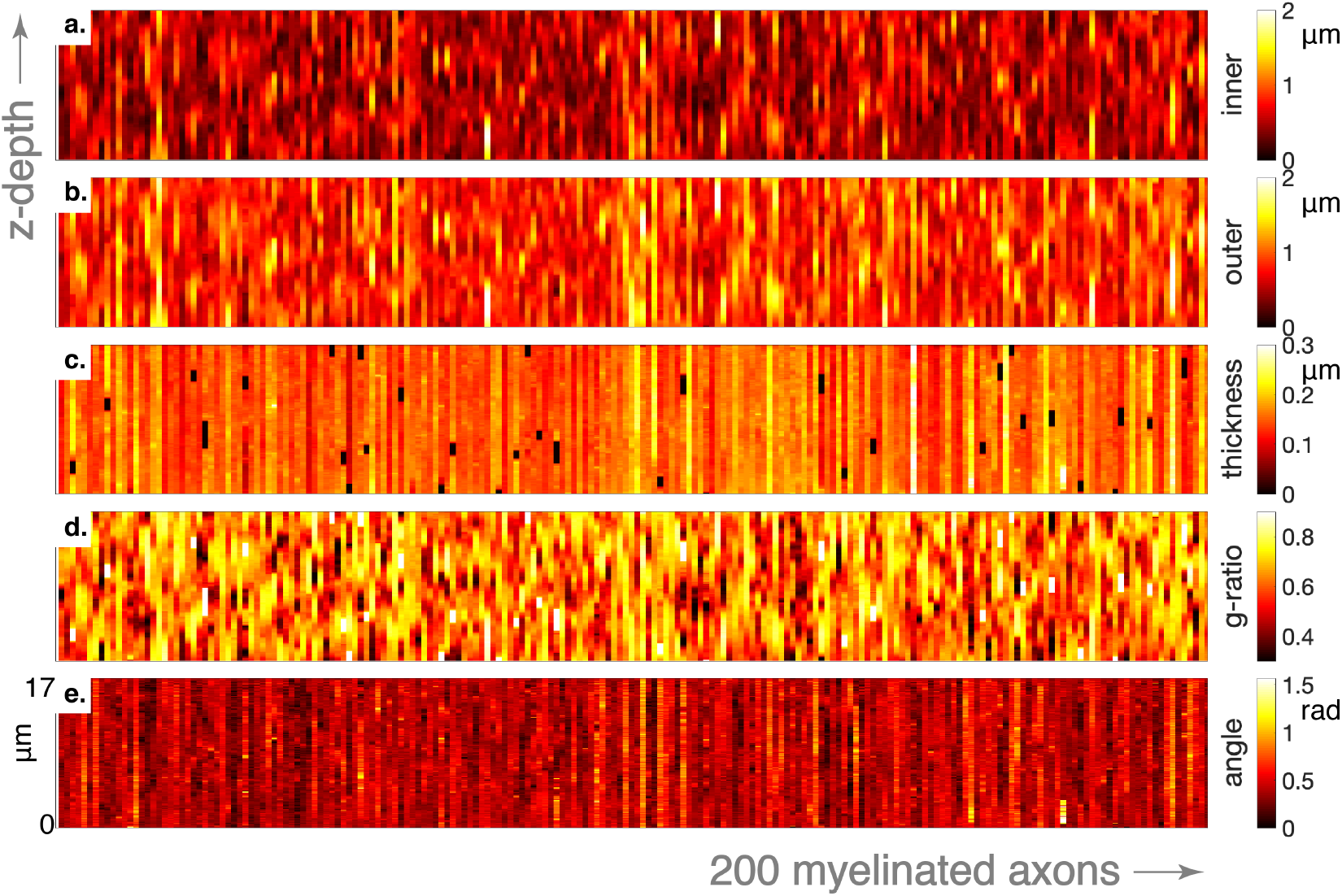
Variations along the axons. The heatmaps show five different features for 200 randomly selected axons over its extent. Each column represents a different axon. a.) Inner diameter of 200 myelinated fibres over ~17 μm of their length. b.) Outer diameter. c.) Myelin thickness is relatively constant within axons (except for the black patches representing nodes of Ranvier). d.) G-ratio variation along axons (nodes of Ranvier are white patches here with g=1). e.) Angle with the bundle’s mean orientation.

**Figure 10.**
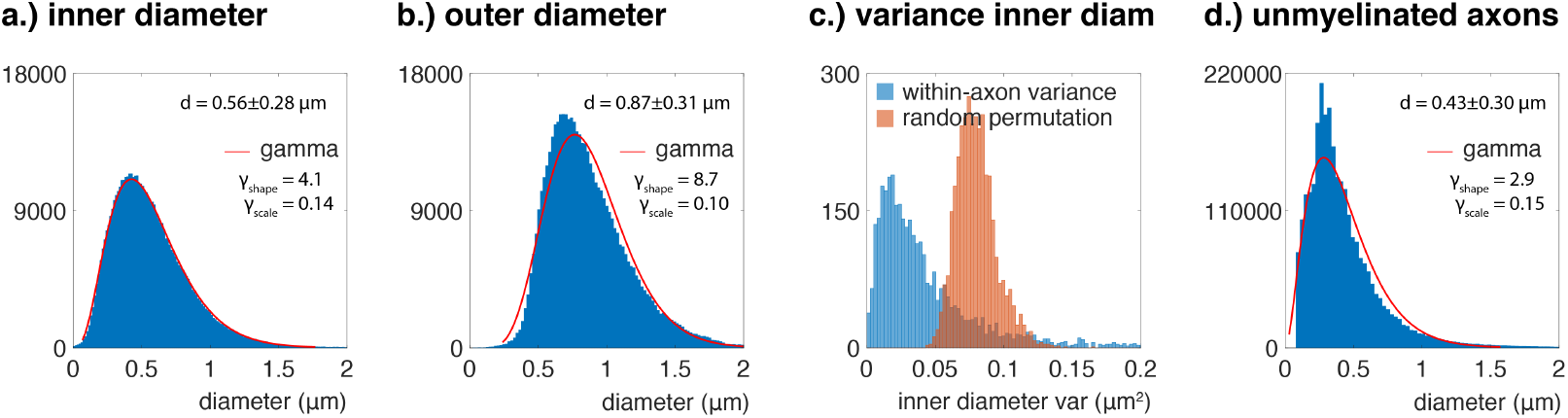
Axon diameter distribution a.) The distribution of equivalent (circle) diameters of the myelinated axons (494,891 cross-sections). b.) The distribution of the equivalent outer diameter of the myelin sheath. c.) Variation in the axon diameter over the axon occurs, but is considerably less than than the variance of a random permutation of the distribution. d.) The distribution of the equivalent diameter of unmyelinated axons.

The myelinated axons had an average equivalent cross-sectional diameter of 0.56 μm (sd 0.28 μm; median 0.51 μm) and their size distribution is well-described by a gamma distribution with shape *k*=4.1 and scale *θ*=0.14 (Figure 10a). The outer diameter, including the myelin sheath, was 0.87 μm (sd 0.31 μm; median 0.81 μm), on average, and deviated from a gamma distribution. For unmyelinated axons (Figure 10d), a mean diameter of 0.43 μm (sd 0.30 μm; median 0.36 μm) was found.

To assess the sources of variance of the axon diameter within and across myelinated axons, we calculated a value for the variance of the equivalent axon diameters of the 2D-labels comprising each axon, as well as from a random permutation of all these 2D cross-sections in the dataset. The variance of the diameter within individual myelinated axons over z-sections is much smaller as compared to when the values of the section are randomly permuted across the dataset (Figure 10c), suggesting that much of the variance in the diameter distribution can be ascribed to axons having a range of calibres. Yet, the variation over sections along the axons is not negligible. Figure 9 shows depth profiles of a set of 200 randomly selected axons from the set that fully traversed the volume over all sections. Variation over depth of the myelinated axon diameter is commonly observed (Figure 9b) and has a typical period of >2 μm and may well have alternations of the inner diameter (Figure 9a) between very thin segments (<0.5 μm) and wide segments (>1 μm).

The g-ratio when measured for each axon and cross-section (Figure 11a) had an average of 0.62 (sd: 0.12; median 0.63). When calculating a per-axon aggregate g-ratio from the axon and myelin volume (Figure 11b), however, the g-ratio mean (and distribution) was 0.67 (sd: 0.079; median 0.67). This variation in g-ratio (Figure 11c) is primarily driven by the axon diameter (Figure 9a) rather than the myelin thickness which is homogeneous over the axon (Figure 9c). This pronounced g-ratio variation over the extent of the axons has the consequence of diverging averages, because the true g-ratio average over sections is lower than the aggregate g-ratio calculated from the volumes pooled across sections as 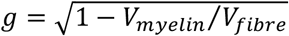.

**Figure 11.**
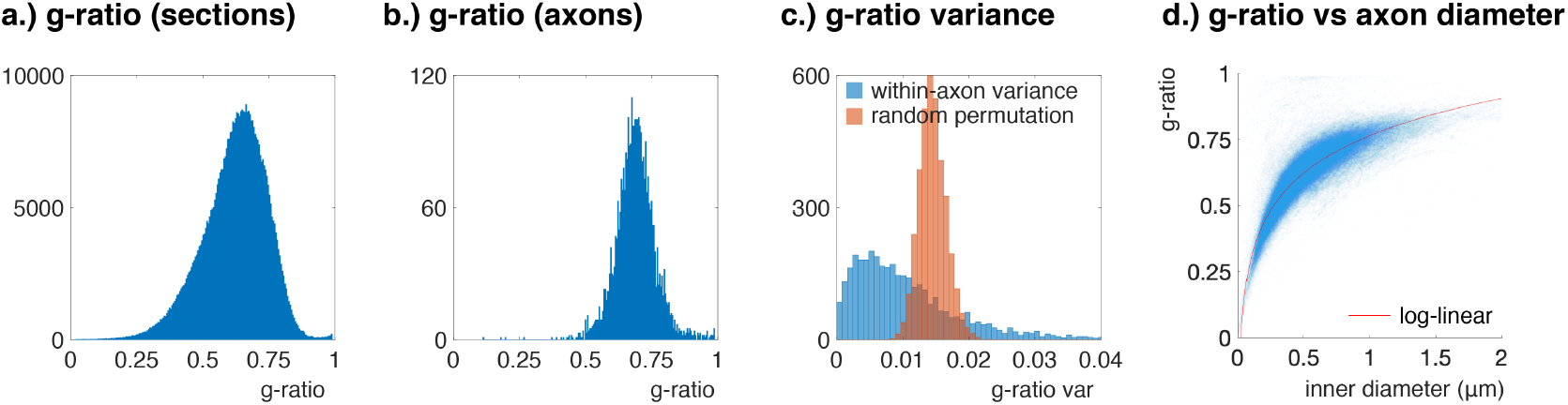
G-ratio distribution. a.) The distribution of g-ratio of all the cross-sectional axons (494,891 cross-sections). b.) The shape of the distribution changes when pooling the g-ratio over the axons, because within-axon g-ratio variation (due to varying axon diameter) is averaged out. E.g. the many thin segments of myelinated axons (that mostly exhibit low g-ratios) may the reason for the skewness of the distribution in a), which is obscured through averaging over the sections of axon. c.) The g-ratio shows variance over the extent of the axon. Differences across axons are larger, yet overall modest as seen from the fairly tight distribution in (b). d.) The relation between the inner axonal diameter and g-ratio can be described by a log-linear fit as proposed in [39].

The dispersion of the myelinated axons is shown in Figure 12. Although the top view on the axons suggests a high dispersion, the side view indicates a relatively homogeneous bundle (Figure 12a). The histogram (Figure 12b) and orientation distribution (Figure 12c) confirm a tight distribution around a mean that is 12° off the z-axis. The dispersion is near-isotropic for this sample with κ_1_ = 23.6 (ODI_1_ = 0.0272) and κ_2_ = 16.7 (ODI_2_ = 0.0381) for a fit to a Bingham distribution [40].

**Figure 12.**
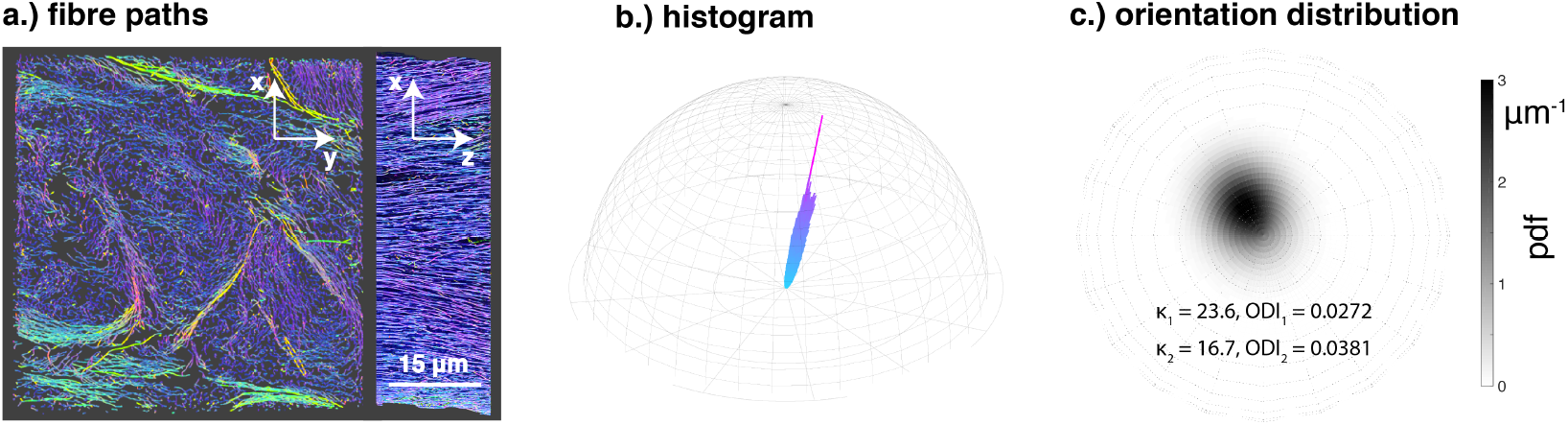
Dispersion a.) Myelinated fibre pathways through the dataset as seen along the section direction (left panel) and in an othogonal view (right panel). b.) Histogram of fibre segments. c.) The fibre orientation distribution plotted on a sphere in a view along the section direction.

## Discussion

In the development of novel methods in MRI research, identification of relevant and informative features in the MRI signal has much to gain from accurate models of the microstructure from which the signal is generated. We have presented a pipeline to derive a representation of the brain’s WM microstructure at a (sub)cellular level by dense segmentation. It includes the main cellular compartments (myelinated axons, unmyelinated axons, glial cells, blood vessels) and various subcellular features of the tissue relevant for a host of physiological processes (myelin sheaths, mitochondria, nodes of Ranvier). This allows detailed interrogation of the datasets for tissue properties and provides a testbed for probing specific microstructure manipulations. The methods as well as data and accompanying segmentations are made available as a resource to the neuroscience community.

We have attempted to design a pipeline that is as comprehensive as possible with regards to the content of the acquired 3D EM data. We segmented the full datasets and provide dense annotation: labelling all voxels as part of a specific cell and attempting to assign them to subcellular structures where possible. We will highlight the performance and utility of a number of these compartment annotations, and discuss some relevant features that are lacking or might be inaccurate in our segmentation. Also, we will provide suggestions on avenues for improvement and application.

### Pipeline features

Our pipeline uses progressive step-by-step compartment annotation and refinement, starting with the easiest-to-segment objects down to more challenging objects. At each stage, the already identified structures are masked from the process, reducing errors in segmenting the more challenging objects. We have built our pipeline upon tools that are accessible and readily available (scikit-image, Ilastik, ITK-SNAP, NeuroProof). Where appropriate, computational cost and the burden of manual intervention was reduced by working on more manageable low-resolution images, with implementation of upsampling methods for representing the segmentations at the full resolution of the acquisition. For other steps (Ilastik pixel classification, generation of supervoxels, NeuroProof agglomeration), high performance distributed computing with trivial parallelization (blockwise processing) to speed up computation or large memory nodes (separation of myelin sheaths) were used.

A relevant novel feature that has been introduced in our segmentation approach is the careful consideration of the accurate segmentation of abutting myelin sheaths. Myelin sheath thickness is a parameter of considerable interest in neuroscience. It is an important factor in signal conduction velocity [41] and may be used as a marker for disease [42]. Together with the axon diameter it determines the g-ratio. Both the aggregate axonal calibre and g-ratio have been suggested to be MRI-detectable [43,44], although questions remain regarding both in-vivo translation and accuracy. In our data, myelin thickness was found to be roughly homogeneous within axons, but neighbouring axons may have very different myelin sheath thickness. Because abutting sheaths cannot be distinguished by textural features in EM, a common approach to separate sheaths is by watershed of the Euclidian distance transform. However, the watershed line on the midway point between the axons is an inaccurate representation of the myelin structure. As the myelin properties are one of the main measures of interest from the segmentation, we found it essential to improve the separation of the sheaths. In our pipeline we used an iterative weighted watershed that takes the median thickness of the sheath over the axon into account. The median thickness yields a good prediction of the true sheath thickness at the site of touching axons, provided the sheath thickness is fairly constant over the individual axon and the largest surface of the axon does not touch other myelinated axons. We have shown that this inaccuracy of a basic watershed on the distance transform affects a large proportion of axons and that the misestimation of the thickness is non-negligible. For accurate quantification of myelin thickness from segmentations using the watershed approach, it is therefore necessary to employ a correction that counters this bias.

### Compartment properties

The diameter of myelinated axons as measured by the equivalent circle diameter in the dataset described here (mean 0.56 μm; sd 0.28 μm) is in accordance with the average diameter reported in similar recent studies by West et al., 2015 [45] (mean 0.56 μm; sd 0.32 μm), Sepehrband et al., 2016 [46] (mean 0.54 μm; sd 0.28 μm) and Abdollahzadeh et al. 2019 [28], but markedly different from Lee et al., 2019 [12] (mean 0.99 μm; sd 0.42 μm). This difference may be explained by selection of larger axons through the random walker segmentation that disregards leaky axons.

In contrast to what was found by Sepherband et al., 2016 [47] and Lee et al., 2019 [12], we did not obtain a better fit of the inner axonal diameters to the generalized extreme value distribution (not shown) as compared to the gamma distribution, although the log-likelihood was marginally better for the generalized extreme value distribution. For the outer diameter, however, the generalized extreme value distribution was markedly better than the gamma distribution. Our measurements are in agreement with Lee et al., 2019 [12] about within-axon variance of the myelinated axon diameter, arriving at slightly higher, but comparable, coefficients of variation (CV_inner_: mean 0.37, median 0.35, sd 0.15; CV_outer_: mean 0.23, median 0.22, sd 0.10).

In this study, we specifically note that the g-ratio not only varies across axons, but also within axons (CV: mean 0.16, median 0.15, sd 0.078) due to pronounced variation of axon diameter while maintaining constant myelin thickness over the axon’s extent (except for nodes of Ranvier, which were excluded from g-ratio analysis;). Although it does not invalidate MRI-based g-ratio models (since they are specifically designed to be agnostic about the internal distribution of myelin within the voxel), this point has been overlooked in the literature.

The dispersion of myelinated axons was low and showed little directional difference. The Bingham distribution fit yielded κ_1_ = 23.6 and κ_2_ = 16.7, where estimates of other studies that evaluate dispersion in 3D are: κ_1_ = 19 and κ_2_ = 5 [12]; κ_1_ = 21 and κ_2_ = 12 [48]. During our acquisition, the dataset location was specifically selected as a region where the top surface of the sample block visible during setup of the acquisition did not contain many cell bodies and blood vessels, but rather a region dominated by axons, with a quasi-circular cross-section. It is probable that this selection bias is responsible for the absence of dispersion in the dataset and that it represents a value near the lower bound of dispersion found in the corpus callosum.

### Limitations

Despite our best efforts to minimize errors in the segmentation, inaccuracies remain. Some of these are due to our methods not functioning as intended, others due to unmet demands (e.g. non-trivial myelin geometry) and yet others are inaccuracies of representation of the compartments (e.g. omission of the extracellular space).

One challenge lies in the complexity of the organization of even the most regularly ordered white matter bundle investigated here. An example is where myelin sheaths do not form simple wraps, but expand from oligodendrocyte process towards multiple axons or are not tightly compacted. These cases are not handled by algorithms in our pipeline. Improvements may be achieved in future implementations by specifically modelling the myelin sheath as continuous closed surfaces.

Our pipeline works best for coherent axon bundles oriented perpendicular to an ordinal axis (the direction of sectioning). Axons traversing the volume obliquely have aberrant cross-sectional shape and labels are more likely to be rejected in our 2D processing steps. For these axons, more extensive manual correction was required which would increase with more heterogeneous orientation distributions.

We have used machine learning classification of the unmyelinated axons by means of the NeuroProof software library. Although this tool delivers an initial dense segmentation, it still requires extensive proofreading to correct split/merge errors. We have not corrected these errors in the datasets presented here, as our intended applications do not strictly necessitate it. If required for the particular application, the unmyelinated compartment could be improved by using proofreading tools such as guided proofreading [49].

Although we have achieved segmentation of cellular constituents of the tissue, it has to be considered that 20% of the volume of white matter tissue is extracellular space (ECS) [50]. The sample preparation procedures in EM reduce the ECS volume to the point that there is little space between axons. This poses a problem for the accurate representation of the tissue state *in vivo*. This is particularly relevant in the application of diffusion MRI simulations, because the protons diffusing unrestrictedly in the ECS can be an important contributor to the diffusion MRI signal. A way to handle the absence of the ECS in the EM data is to artificially erode the individual axon labels until they occupy a volume fraction of 0.8, after which the aggregate voxelvolume can be rescaled to reinstate the original axon volumes. However, it is not certain that the shape of the artificially induced ECS is a good representation of the *in vivo* situation. We have explored alternative EM preparation methods that preserve the ECS [21], but it is still not known if the ECS morphology in these preparations is representative of *in vivo* tissue structure. This specific issue is just one example of the general concern of morphological changes of the cells associated with various preparation protocols. For instance, the cross-sectional shape of axons is affected by artefacts from chemical fixation and ethanol dehydration, as compared to their shape observed following high pressure freezing and freeze substitution [51]. In sum, tissue preparation methods may be an important caveat in the interpretation of morphological shape measures and absolute volumetric measurement. Therefore, proper consideration must be given to any conclusions derived from these measures when generalising them to the *in vivo* situation.

## Conclusions

We have presented an approach for dense and detailed segmentation of 3D EM data of WM. A novel element in the pipeline of specific interest for white matter investigations is the method for myelin segmentation that yields accurate boundaries for the individual axons. The segmentation consists of individual cells in the volume, as well as nested subcellular components, such as myelin, mitochondria and the nodes of Ranvier. These objects can be interrogated for their morphological properties and can be used in validation and development of biophysical models for predicting MRI signals. This work has presented benchmark statistics of MRI-accessible microstructural properties (axon diameter and g-ratios). Future work will focus on the comparison between white matter regions and application of models in in silico experiments.

## Acknowledgements

The research was funded by a Wellcome Trust Senior Research Fellowship (202788/Z/16/Z). *The Wellcome Centre for Integrative Neuroimaging is supported by core funding from the Wellcome Trust (203139/Z/16/Z). Electron microscopy was conducted at the Dunn School EM Facility.* The BBSRC provided Advanced Life Sciences Research Technology Initiative 13 funding for SBF-SEM through Grant BB/C014122/1 (to Chris Hawes).

