## Supplementary Figure 1 for "A semi-automated approach to dense segmentation of 3D white matter electron microscopy"

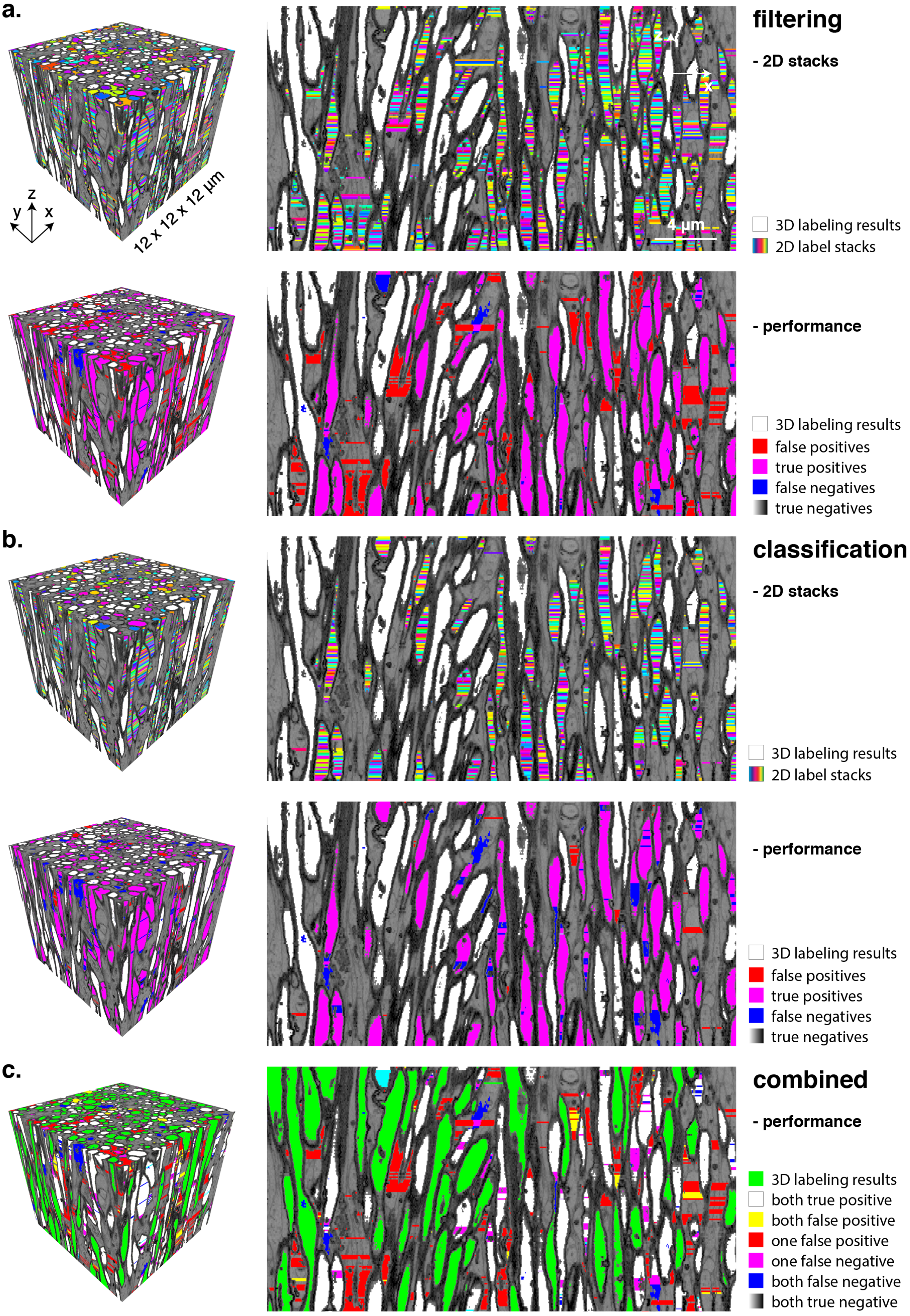


**Supplemtary Figure 1. Method comparison for 2D label selection/rejection. a.) The selected 2D labels of the feature filtering method (top) contain a large number of false positive (bottom; red), while performing well on false negatives (bottom, blue). b.) The stacks of labels obtained with the feature classification method (top) do not include many false positives (bottom, red), although the number of false negatives are increased in comparison to the feature filtering method (bottom, blue). c.) Overlay of the FP/TP/FN/TN analysis of both methods.**
