## Supplementary Figure 2 for "A semi-automated approach to dense segmentation of 3D white matter electron microscopy"

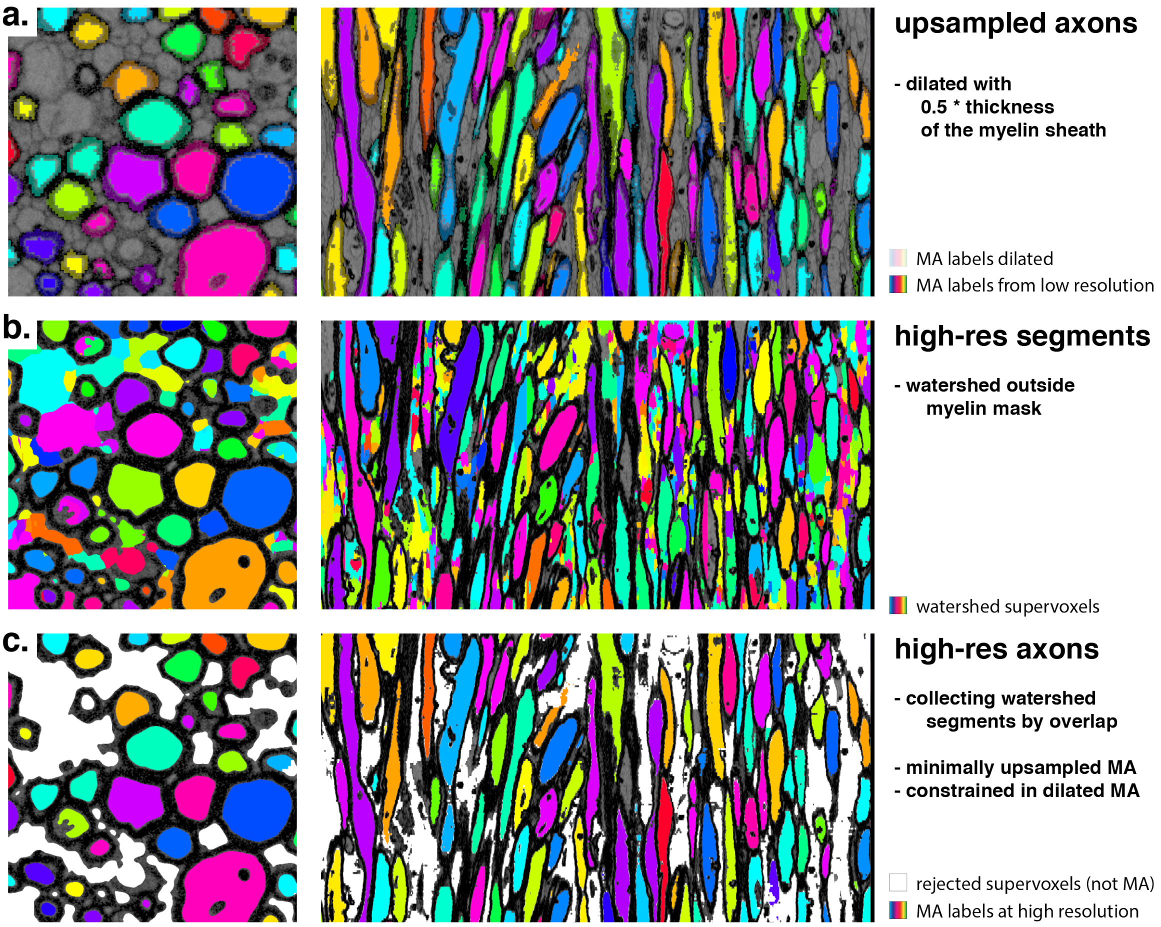


**Supplementary Figure 2. Myelinated axons (MA) at full resolution. a.) upsampled myelinated axon labels and their dilation with half the myelin sheath thickness define the inner and outer boundary of the MA at high resolution. b.) watershed segments from the full resolution data. c.) aggregation of the watershed segments yields MA labels at high resolution, discarding segments of the unmyelinated axon space (in white).**
